# Identification of key tissue-specific, biological processes by integrating enhancer information in maize gene regulatory networks

**DOI:** 10.1101/2020.06.16.155481

**Authors:** Maud Fagny, Marieke Lydia Kuijjer, Maike Stam, Johann Joets, Olivier Turc, Julien Rozière, Stéphanie Pateyron, Anthony Venon, Clémentine Vitte

## Abstract

Enhancers are important regulators of gene expression during numerous crucial processes including tissue differentiation across development. In plants, their recent molecular characterization revealed their capacity to activate the expression of several target genes through the binding of transcription factors. Nevertheless, identifying these target genes at a genome-wide level remains a challenge, in particular in species with large genomes, where enhancers and target genes can be hundreds of kilobases away. Therefore, the contribution of enhancers to regulatory network is still poorly understood in plants. In this study, we investigate the enhancer-driven regulatory network of two maize tissues at different stages: leaves at seedling stage and husks (bracts) at flowering. Using a systems biology approach, we integrate genomic, epigenomic and transcriptomic data to model the regulatory relationship between transcription factors and their potential target genes. We identify regulatory modules specific to husk and V2-IST, and show that they are involved in distinct functions related to the biology of each tissue. We evidence enhancers exhibiting binding sites for two distinct transcription factor families (DOF and AP2/ERF) that drive the tissue-specificity of gene expression in seedling immature leaf and husk. Analysis of the corresponding enhancer sequences reveals that two different transposable element families (TIR transposon *Mutator* and MITE *Pif/Harbinger*) have shaped the regulatory network in each tissue, and that MITEs have provided new transcription factor binding sites that are involved in husk tissue-specificity.

**Significance:** Enhancers play a major role in regulating tissue-specific gene expression in higher eukaryotes, including angiosperms. While molecular characterization of enhancers has improved over the past years, identifying their target genes at the genome-wide scale remains challenging. Here, we integrate genomic, epigenomic and transcriptomic data to decipher the tissue-specific gene regulatory network controlled by enhancers at two different stages of maize leaf development. Using a systems biology approach, we identify transcription factor families regulating gene tissue-specific expression in husk and seedling leaves, and characterize the enhancers likely to be involved. We show that a large part of maize enhancers is derived from transposable elements, which can provide novel transcription factor binding sites crucial to the regulation of tissue-specific biological functions.

## Introduction

Enhancers are key regulators of the spatio-temporal expression of genes in eukaryotes, in particular during development [1, 2]. Their regulatory effect is mediated by the binding of transcription factors (TFs), which interact with target gene promoters through 3D-loops over distances reaching several dozens of megabases in some species [3, 4]. The binding of a single TF is often not sufficient to activate the expression of a gene, and generally several TFs act together to increase or decrease the regulatory potential of a given enhancer [1]. Groups of enhancers characterized by similar content in transcription factor binding sites (TFBSs) have been shown to co-regulate genes involved in the same biological pathways, thus shaping a complex regulatory network controlling the tissue-specific expression of genes involved in particular biological functions [5, 6]. While enhancers have been identified as key players in the wiring of the developmental gene regulatory network in mammals [7, 5], this question remains largely unexplored in plants [2].

Recent combined analyses of DNA methylation, chromatin accessibility and histone marks have led to the genome-wide characterization of thousands of putative active enhancers in plants [8, 4, 9, 10, 11, 12, 13]. Distance between each enhancer and its nearest gene varies strongly depending on the species, ranging from about 2 kb in *Arabidopsis thaliana* to 1 Mb in barley, and is largely correlated to genome size [9]. In maize (*Zea mays* ssp. mays), 3D chromatin folding analyses showed that about 25%-40% of enhancers are not targeting their closest gene, and that 34% of enhancers potentially regulate several genes [4, 14]. These results highlight the difficulty to identify the regulatory relationships between enhancers and their target genes in plants with large genomes.

How enhancers arise and rewire the gene regulatory network in plants is unclear. Transposable Elements (TEs) of various superfamilies have been proposed as a source of new regulatory elements [15] and have been shown to be involved in the rewiring of gene regulatory networks for some key tissue-specific biological functions in animals [16]. In plants, examples of enhancers derived from a particular TE have been described [17, 18, 19], and a more general contribution of TEs to *cis*-regulatory elements has been highlighted in some species such as *Capsella grandiflora* [20] and maize, where at least a quarter of the thousands of putative enhancers were found to overlap TE annotations [8, 10]. TEs influencing the response of nearby genes to abiotic stresses have also been described, for example in maize seedlings [21], hinting for an important role of TEs in regulating the expression of genes involved in specific biological functions in plants. Nevertheless, whether TEs contribute to the emergence of tissue-specific gene regulatory networks in plants remains to be fully elucidated.

As of today, it remains both time-consuming and expensive to test the enhancer-target gene regulatory relationship at the genome-wide level using molecular biology approaches such as enhancer reporter assays and CRISPR-Cas manipulation. By offering approaches to model *in silico* the regulatory relationships between heterogeneous components such as TFs and genes, systems biology provides a powerful and cost-effective alternative. Classical co-expression networks allow to group TFs with potential non-TF target genes. However, they do not provide information about whether these regulatory relationships are actually possible in terms of binding of the TF to regulatory elements associated with the target genes. Integrating information about the genes *cis* -regulatory sequences, in particular about which TFBS they harbor, allows to connect TFs more directly to their potential target genes. This information can then be integrated with gene co-expression information to generate bipartite TFs-genes networks. Such systems biology approaches have contributed to decipher the role of promoter-binding TFs in the regulation of their target genes in tissues or cell cycle stages in fungi and animals [22, 23, 24, 25], and to identify the impact of disease on the wiring of tissue-specific regulatory networks in humans [26]. With recent advances in active enhancer characterization, TFBS annotation, and the generation of expression data from a large number of tissues, these systems biology approaches can now be used in plants and open new opportunities to study the regulatory role of enhancers during plant development. Improvement of genome sequences and TE annotation also allows for characterizing the part of enhancers driven by TEs, and therefore to investigate the potential role of TE sequences in rewiring gene regulatory networks in plants.

In this study, we investigate the interconnection between TFs, enhancers and target genes in maize tissue-specific gene regulation, by comparing the regulatory networks of two types of maize leaves at different developmental stages: immature leaves at V2 seedling stage and husks (bracts) at flowering. Taking advantage of enhancers that were previously predicted by Oka and colleagues to be active in these two organs [8], we analyze their TFBS composition. Using bipartite networks, we then integrate this information with relative genomic position of genes and enhancers together with transcriptomic data, to reconstruct tissue-specific TFs-genes regulatory networks. We identify key TFs that co-regulate groups of genes involved in biological functions crucial for tissue identity, and link these genes to the enhancers that regulate their expression. By analyzing sequences of these enhancers, we show that TIR transposon *Mutator* and MITE *Pif/Harbinger* families are involved in the tissue-specific expression in immature seedling leaf and husk, respectively. We also discover that MITEs harbor conserved sequences that are likely maize-specific TFBS, thus highlighting that TEs are important players in shaping regulatory networks in this species. An online queryable version of the networks is available at https://maud-fagny.shinyapps.io/TF-gene_network_Maize/.

## Results

### Husk and V2-IST-speciflc enhancers are enriched in binding sites targeted by different TF families

We first aimed to characterize the TFBS content of active enhancers in husk and V2-IST. To this end, we extracted sequences of the 1495 putative active enhancers (hereafter called enhancers) obtained from Oka and colleagues, among which 1097 were found specifically active in husk, 175 specifically active in V2-IST, and 223 active in both tissues. We *in silico* annotated the TFBSs located in these enhancers by scanning for known plant TFBSs (Figure 1). After selecting for TFBSs with a Benjamini-Hochberg-corrected *p*-value below 0.01 (see Materials and Methods), we retained 18,348 TFBSs corresponding to 62 transcription factors. Among all enhancers, 524 (34.6%) did not harbor any significant TFBS, including 441 (40.2%) husk-specific enhancers, 34 (19.4%) V2-IST-specific enhancers and 49 (22.0%) shared enhancers. Using a resampling approach (see Materials and Methods), we tested for TFBS enrichment in enhancers of each tissue. We found that 60.6% of the V2-IST-specific enhancers, 60.6% of the husk-specific and 66.8% of the shared enhancers were significantly enriched for TFBSs compared to randomized sequences. On average, an enhancer contained 11.4 TFBSs (ranging from 0 to 255), which covered on average a total of 34.3 bp (ranging from 0 to 442 bp) or 2.5% of the enhancer sequence length.

**Figure 1:**
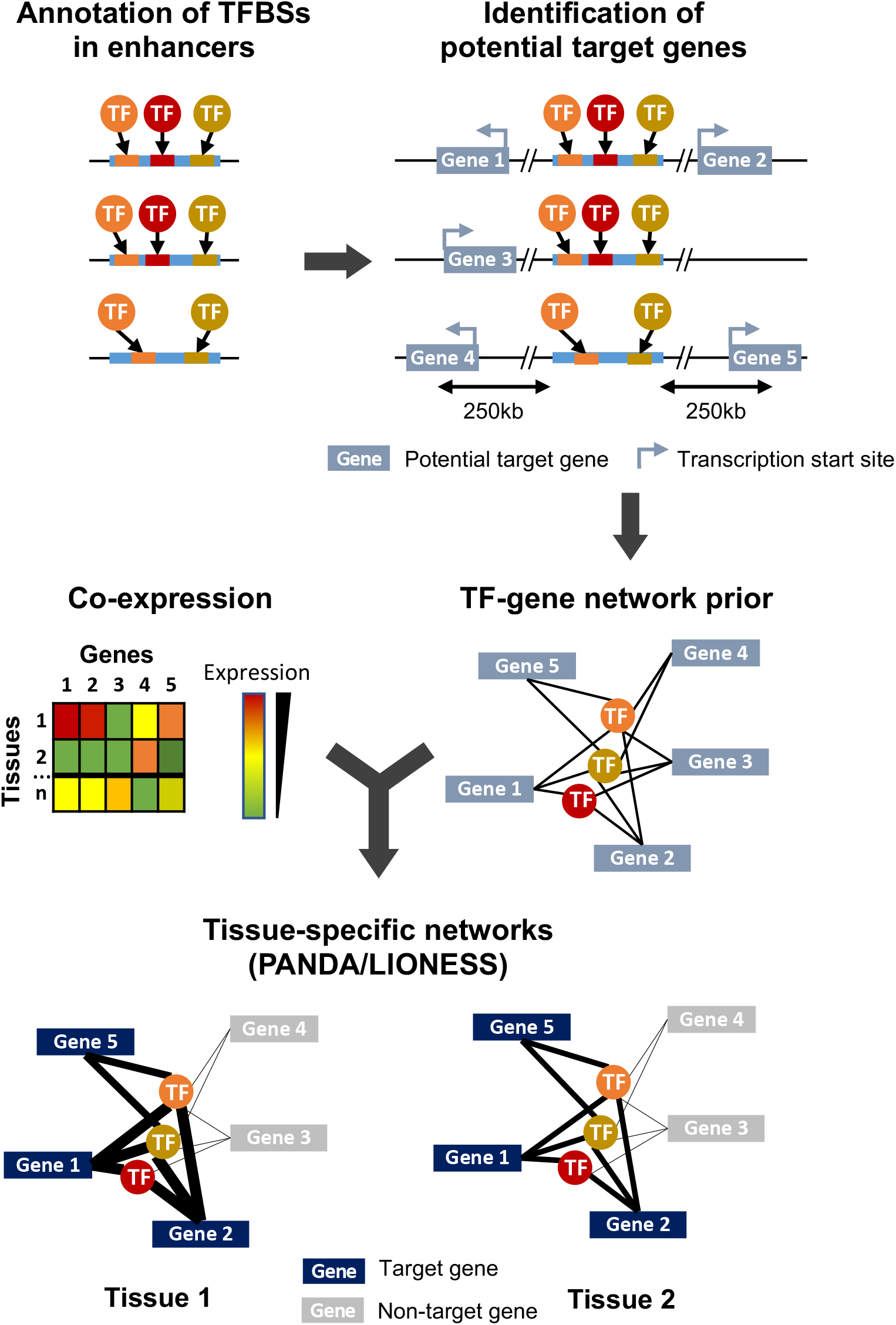
Overview of the study design and methodology. Enhancer sequences are first searched for TFBS motifs using FIMO. All genes whose transcription start site is within 250 kb of an enhancer are considered as potential targets, and combination of this genomic information and TFBS annotation leads to the generation of a TFs-genes prior network. In parallel, a gene-gene co-expression matrix is generated from expression data obtained from different samples (here, corresponding to different tissues) and integrated to the TFs-genes prior using PANDA and LIONESS to obtain tissue-specific regulatory networks.

We then compared the TFBS content of enhancers active in husk and these active in V2-IST. Husk enhancers were enriched for binding sites corresponding to 15 TFs, mostly from the C2C2-DOF family (7), but also from the AP2/ERF (5), HD-ZIP (2) and bHLH (1) families. More precisely, in addition to the 3 TFs of the C2C2-DOF family that had significantly more binding sites in the husk enhancers, 4 were found only in husk enhancers (Benjamini-Hochberg adjusted *p*-value lower than 0.05, Supplementary Table S1). TFs from the C2C2-DOF family are known to be mainly involved in response to abiotic and biotic stresses, and are expressed in growing and mature leaves [27, 28, 29]. In contrast, V2-IST enhancers were enriched for binding sites recognized by TFs from the AP2/ERF family, with 17 TFs having significantly more frequent TFBSs in V2-IST enhancers than in husk enhancers (Benjamini-Hochberg adjusted *p*-value below 0.05, Supplementary Table S1). AP2/ERF TFs are known to be involved in plant development and growth, and transition to flowering [30], and are known to be expressed in seedlings [28, 29].

### TFs-genes regulatory interactions recapitulate tissue-specific enhancer activity

To investigate the tissue-specific regulatory relationship between TFs and their target genes, we first built a prior network. In this prior, we considered that a gene was potentially targeted by a TF if an enhancer containing the TFBS recognized by this TF was located within 250 kb of the transcription start site of a gene (Figure 1). All enhancers (*i.e*. these found in husk or in V2-IST) were used. Because about 25% of enhancers are indicated to be downstream of their target genes [4], we included all genes independently of their genomic orientation. To build tissue-specific regulatory networks, we combined this prior with gene co-expression data [22, 25]. We had access to mRNA-seq data from husk and V2-IST (6 replicates each) from Oka *et al*., 2017 [8]. To obtain high confidence tissue-specific networks [25], we enriched this dataset with mRNA-seq data that we generated from 11 tissues in triplicates (see Materials and Methods and Supplementary Table S2). In total, our merged dataset included 45 samples and a total of 46,430 genes. We retained genes expressed in at least 3 samples in at least 1 tissue (see Materials and Methods), corresponding to a total of 36,041 genes. Read counts were then normalized and corrected for the single-end/paired-end mRNA-seq data type (see Materials and Methods). As shown by correlation analyses and principal component analysis (PCA), the data from both datasets are comparable (Supplementary Figure S1). In particular, expression levels from the V2 inner immature tissues from both datasets are strongly correlated (average pairwise correlation coefficient R=0.93), while the average inter-tissue R is 0.83 and the average biological replicate R is 0.95. V2 inner tissues from the two data sets (V2-IST and V2_LFI) also cluster together in the PCA (Supplementary Figure S1).

Out of the 36,041 genes, we identified a total of 8,054 potential target genes that were located within 250 kb of one of the 1,495 enhancers, and that we included in the prior. Among those, 6,459 (80%) were potentially targeted by a single enhancer, 1,269 (16%) by two enhancers and 326 (4%) were potentially targeted by three enhancers or more. A total of 971 enhancers had potential target genes within 250 kb, and each of these enhancers had an average of 10 potential target genes (ranging from 1 to 42). Of the 62 TFs for which TFBSs were identified within the enhancers, 10 were not expressed in any of our samples and were therefore filtered out. Our prior gene regulatory network thus contained 52 TFs that had TFBS in one of the 971 enhancers, and 8,054 genes. Each gene was connected to an average of 8.6 TFs (ranging from 1 to 33), and each TF was linked to an average of 1310 genes (ranging from 2 to 3431).

We combined the prior gene regulatory network to the co-expression matrix obtained from the mRNA-seq normalized data from the 45 samples (13 tissues). We then built sample-specific gene regulatory networks using PANDA and LIONESS [22, 25], and obtained 45 sample-specific TFs-genes regulatory networks (Figure 1). We generated a 2D representation of the sample-specific networks using a uniform manifold approximation and projection for dimension reduction (UMAP) approach (Supplementary Figure S2), and compared it to the PCA results on gene expression data. As expected, samples from the same tissue cluster together on the UMAP, and the V2-specific regulatory networks generated from V2 growing leaves from the formerly and newly generated datasets cluster together. Notably, while in the PCA husk samples were isolated and located close to silk and internode tissues (Supplementary Figure S1), in the UMAP the husk-specific regulatory network was clustered with regulatory networks of other types of mature leaves, thus indicating stronger similarity in terms of gene regulatory network than gene expression levels between tissues that share similar developmental stages.

We sought to validate our approach by examining the tissue-specific regulatory relationships between genes and TFs for textbook cases. Among the three known enhancers included in the study of Oka and colleagues, two had known target genes mapped within 250 kb in the AGPv4 maize genome assembly: these of *tb1* and *bx1* (also known as DICE). The third one, *b1* enhancer has not been assembled in AGPv4, and as such could not be included in our analysis. We found that the regulatory relationship between *tb1* and the TFs binding its enhancer is stronger in husk than in V2-IST (Supplementary Figure S3A). This is in accordance with former observations that the *tb1* enhancer is active in husk but not in V2-IST [8]. In contrast, the regulatory relationship between *bx1* and the TFs binding its enhancer (DICE) were of similar strength in both husk and V2-IST (Supplementary Figure S3B), in accordance with the fact that DICE was shown to be active in both tissues. Hence, our approach, which uses a generic TFs-genes prior common for both husk and V2-IST tissues and co-expression data, is able to retrieve the activated/unactivated states of known enhancers in each tissue specifically.

### Tissue-specific regulatory modules highlight different biological functions in husk and V2-IST

Our first aim was to identify and biologically characterize regulatory networks that were differentially regulated between husk and V2-IST. To this end, we first performed a differential targeting analysis of the V2-IST and husk tissues by comparing the edge weights of the sample-specific networks between the two tissues (see Materials and Methods). We thus identified 2,075 genes that were more highly targeted by TFs in husk and 2,123 genes that were more highly targeted in V2-IST (Benjamini-Hochberg corrected *p*-value of 0.05). The 3,856 remaining genes were not significantly differentially targeted in any of the two tissues. Using a Gene Ontology enrichment analysis, we found that genes with a higher TF-gene regulatory relationship in husk were enriched for biological processes such as “protein phosphorylation”, “cell growth”, and “regulation of intracellular signal transmission” (GO:0006468, GO:0016049, GO:1902531, Supplementary Table S3). In contrast, genes with higher TF-gene regulatory relationship in the V2-IST network were enriched in biological processes related to “cell proliferation”, “regulation of DNA replication”, “regulation of meristem development” and “chloroplast organisation” (GO:0006275, GO:0008283, GO:0048509 and GO:0009658, Supplementary Table S4), reflecting the fact that V2-IST is a growing tissue and contains the apical meristem, while husk is a more mature leaf tissue.

To get further insights into the differential regulation of these two tissues, we then sought to identify and biologically characterize tissue-specific TFs-genes regulatory modules. To this end, we obtained two tissue-specific networks, one for husk and one for V2-IST (see Materials and Methods) and compared their structure using ALPACA (see Materials and Methods) [26]. This allowed us to identify regulatory modules (*i.e*., groups of TFs that were co-regulating groups of genes) in each tissue-specific network and to compare the modules between husk and V2-IST. We identified 71 modules in the V2-IST-specific network, and 67 modules in the husk-specific network. Among them, respectively 12 and 11 modules contained at least one TF and five genes, and were retained for further investigation. In order to identify shared and tissue-specific modules, we then compared their gene content between husk and V2-IST using the jaccard index. Nine of them had high jaccard indexes (greater than 0.5), indicating that they were very similar between the husk and V2-IST networks (grey modules in Figure 2A and B and Supplementary Table S5). They included 66.8% of the 8,054 genes included in the network prior and contained 570 genes on average (ranging from 5 to 1269 genes). Gene Ontology enrichment analyses showed that the genes contained in these modules are involved in basic biological functions such as protein metabolism, macromolecular complex organisation, and defense response, which are expected to be shared between the two tissues (Supplementary Table S6).

**Figure 2:**
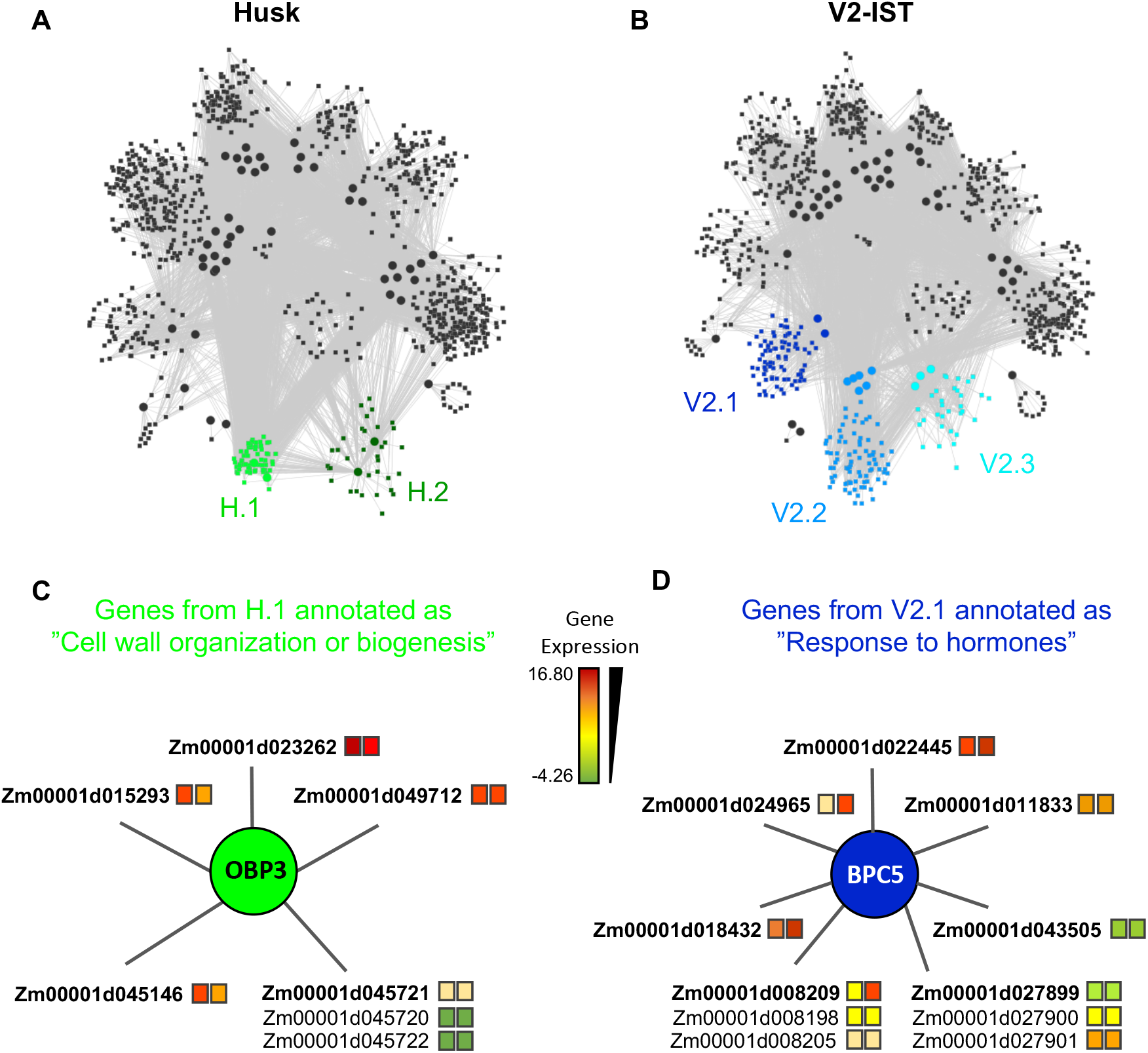
Tissue specific modules. **A.** Module structure of the husk-specific network. The two husk-specific modules (H.1 and H.2) are highlighted in green. **B.** Module structure of the V2-IST-specific network. The three V2-IST-specific modules (V2.1, V2.2 and V2.3) are highlighted in blue. **A-B.** Modules in grey are shared between the husk-specific and the V2-IST specific networks. **C.** Detailed view of the subset of the husk-specific H.1 module that contains the genes annotated as “cell wall organization or biogenesis”. **D.** Detailed view of the subset of the v2-IST-specific V2.1 module that contains the genes annotated as “response to hormones”. **C-D** Colors of the squares located right to gene IDs indicate average expression levels in husk (left square) and V2-IST (right square). When several genes are potentially targeted by the same enhancer, they are represented with a common edge, and the top target is ranked first and highlighted in bold. The TFs that regulate the genes are represented as circles. Because TFBS annotation arise from *Arabidopsis thaliana*, names of TFs are these of this species. The maize ortholog of *obp3* is *dof27*, that of *bpc5* is *bbr4*. Similar information can be retrieved for all genes of the module using the R application we developed https://maud-fagny.shinyapps.io/TF-gene_network_Maize/.

Besides these nine shared regulatory modules, we found five other modules, which were tissue-specific and included two husk-specific modules (containing 61 and 349 genes, respectively) and three V2-IST-specific modules (containing 319, 811, and 859 genes respectively, Supplementary Table S5 and Figure 2A and B). These modules tended to be smaller than the shared ones. We next performed a Gene Ontology enrichment analysis on the genes contained in each tissue-specific module, and focused on GO biological processes that were only significant in tissue-specific modules.

Among the two husk-specific modules, the largest one (349 genes, H.1 in Figure 2A) is enriched for genes involved in “cell wall organization or biogenesis” (GO:0071554, elim algorithm from topGO *p* = 0.009, Supplementary Table S7). This module is clustered around OBP3 (ortholog of maize DOF27), a C2C2-DOF TF known to be involved in leaf development and light signalling in maize mature leaves. Its TFBS is enriched within enhancers activated in husk as compared to V2-IST (Supplementary Table S1). It notably regulates *Zm00001d015293* (Figure 2C), a top target of the husk-specific H535 enhancer (Figure 3) known to be involved in leaf development in rice [31] and to be expressed at the basis of mature leaves [28, 29]. Other interesting targets are *Zm00001d023262* (*brick3*), and *Zm00001d045720/Zm00001d045721/Zm,00001d045722*, three genes coding for proteins of the TBL family, which is involved in trichome morphogenesis and secondary cell wall morphogenesis (Figure 2C) [32]. Most of OBP3 target genes are more expressed in husk than V2-IST, as shown in Figure 2B.

**Figure 3:**
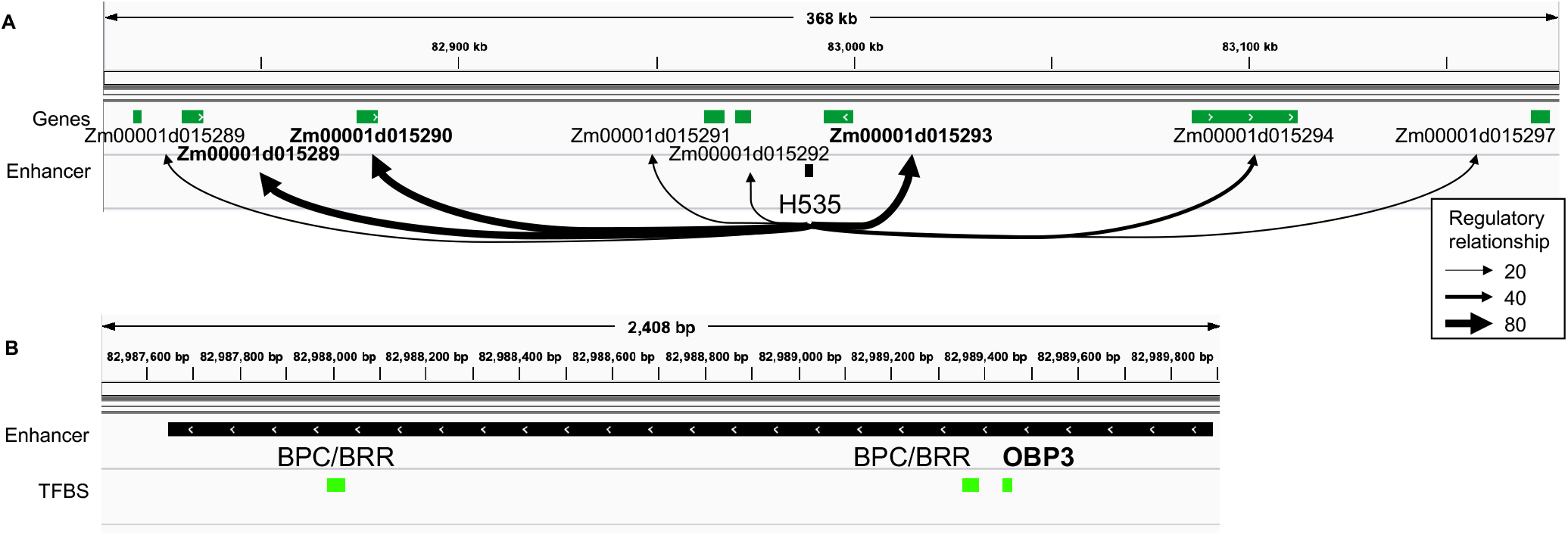
Identification of top targets of a husk-specific enhancer: example of enhancer H535. **A.** Representation of the regulatory relationships between H535 and all of its potential target genes. Thickness of arrows represents average targeting (regulatory relationship) of the genes by all the TFs potentially binding the enhancer. H535 top target genes are highlighted in bold. **B.** TFBS content of the H535 enhancer. OBP3 is highlighted in bold, as it is articulating the “heterocyclic compound binding” husk-specific regulatory module that contains Zm00001d015293.

The small husk-specific module of 61 genes was particularly interesting (H.2 in Figure 2A). It is enriched in genes involved in the molecular function “heterocyclic compound binding” and “organic cyclic compound binding” (GO:1901363 and GO:0097159, elim algorithm from topGO *p* = 0.008, Supplementary Table S7), and it is centered mainly around AT1G12630, a TF from the AP2/EREB family and also includes ERF38, a TF from the DREB subfamily A-4 of the AP2/ERF family (Supplementary Figure S4A). Both *erf38* maize ortholog *erf039* and *At1g12630* maize ortholog *ereb10* are over-expressed in husk as compared to V2-IST (log2 fold changes of 2.4 and 0.9, respectively, and Mann-Whitney U test *p*-value of 2.2 × 10^−3^ and 4.1 × 10^−2^, respectively). The expression levels of both of these TFs positively correlate with the expression level of their target genes involved in “heterocyclic compound binding” across all 45 samples and 13 tissues (Supplementary Figure S5 and Supplementary Figure S6). Accordingly, most of these target genes are over-expressed in husk as compared to V2-IST (Supplementary Figure S4A). Finally, ERF38 TFBSs are only found in enhancers that are active in husk but not in V2-IST (Supplementary Table S1), thus highlighting the importance of this TF in husk-specific gene expression regulation.

The largest V2-IST-specific regulatory module (859 genes, V2.1 in Figure 2B) is enriched in genes involved in several biological processes (Supplementary Table S7) including “sulfur compound biosynthetic process” (GO:0044272, elim algorithm from topGO *p* = 0.002), “regulation of nucleic acid-templated transcription” (GO:1903506, elim algorithm from topGO *p* = 0.004) and “response to hormones” (GO:0009725, elim algorithm from topGO *p* = 3.9 × 10^−2^). In this module, genes classified under the “response to hormones” gene ontology are regulated by BPC5, a member of the BBR/BPC TF family involved in Polycomb complex recruitment (Figure 2D). Consequently, enhancers carrying BPC5 TFBSs are inactive when *bpc5* is expressed and active otherwise. Accordingly, *bpc5* maize ortholog, *bbr4* is less expressed in V2-IST than in husk (log2 fold change −0.4 and Mann-Whitney U test *p* = 2.2 × 10^−3^), its expression is anti-correlated with most of its target genes (Supplementary Figure S7), and its target genes are generally more expressed in V2-IST than in husk (Figure 2C). Notably, its targets include *Zm00001d043505*, a phosphotransmitter [33] known to be involved in the response to cytokinin, a hormone promoting cell division (Figure 2D). *Zm00001d008209* is encoding a protein of the cyclophilin/peptidyl-prolyl cis-trans isomerase family. This gene family is strongly expressed in seedling and growing tissues and is involved in regulating maize development [34].

The second largest V2-IST-specific module (811 genes, V2.2 in Figure 2B) is enriched for genes involved in “cellular macromolecule biosynthetic processes” (GO:0034645, elim algorithm from topGO *p* = 5.0 × 10^−3^, see Supplementary Figure S4B and Supplementary Table S7). Finally, the smallest V2-IST-specific module (319 genes, V2.3 in Figure 2B) is enriched for genes involved in “protein complex assembly” (GO:0006461, elim algorithm from topGO *p* = 7.0 × 10^−3^, see Supplementary Figure S4C and Supplementary Table S7). Both modules are clustered around TFs of the AP2-ERF family including RAP2-12 for the first one (Supplementary Figure S4B) and ERF5 and ADOF1 for the second one (Supplementary Figure S4C). The corresponding TFBSs are enriched in enhancers activated in V2-IST compared to those activated in husk. Altogether, these results show that the TF-gene regulatory networks that we reconstructed allow to identify tissue-specific regulatory modules whose functions are in agreement with the tissue analyzed, as well as key TFs and their target genes underlying these functions.

### Transposable elements are a source of TFBS sequences in tissue-specific enhancers

We took advantage of the regulatory networks and modules we identified to investigate the role of transposable elements (TEs) in the tissue-specific regulation of gene expression. We first compared the TE sequence content of husk-specific and V2-IST-specific enhancers. To this end, we annotated TE in enhancers using a recently updated TE database [35]. We found that of the 971 enhancers present in the prior, 555 (57.2%) were included in, or partially overlapping at least one TE. On average, when an enhancer overlapped a TE, about 18.7% of the enhancer sequence was covered by the corresponding TE sequence (ranging from 0.4% to 100%). We then tested the relative enrichment in TE superfamilies of husk-specific and V2-IST-specific enhancers (Figure 4A). Because husk-specific enhancers were significantly closer to their nearest genes than V2-IST specific and shared enhancers (average 25,722 bp and 32,075 bp, respectively - Mann-Withney U test one-sided *p* =7.3 × 10^−3^) and the distribution of TE families is strongly affected by distance to the closest gene, we used randomly chosen genomic sequences on the same chromosome, with the same size and distance to the closest gene than the enhancers to build *χ*^2^ expected null distributions for each enrichment test (see Materials and Methods). We then compared the *χ*^2^ obtained using the real enhancers with the expected null distribution. We found that husk-specific enhancers are enriched in miniature inverted-repeat transposable elements (MITEs, a group of non-autonomous DNA transposons) as compared to the V2-IST and shared enhancers (odds ratio of 1.8, resampled *χ*^2^ *p* = 0.05).

**Figure 4:**
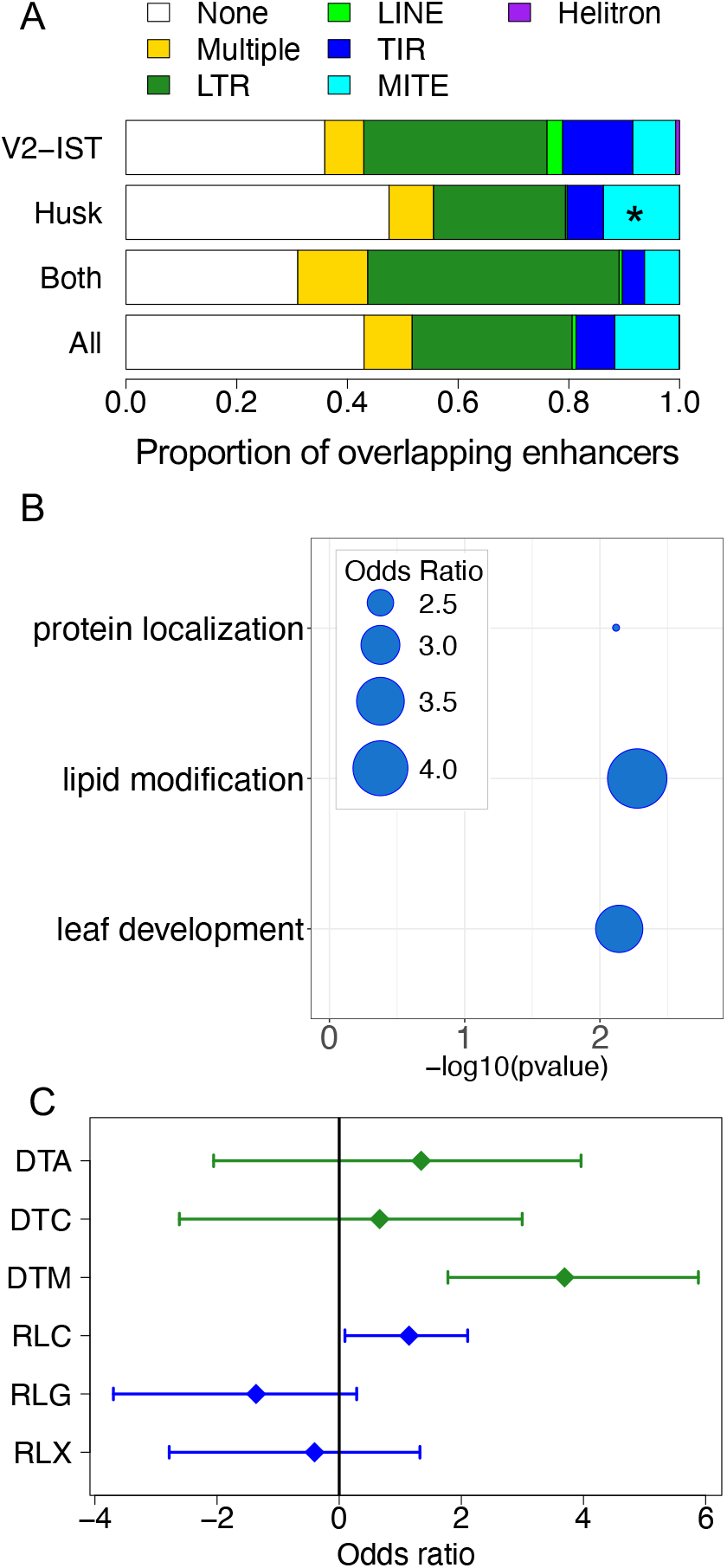
The role of transposable elements in tissue-specific gene expression regulation. **A.** Proportion of different TE orders in enhancers. “None” corresponds to absence of overlap with a TE, and “Multiple” corresponds to presence of overlaps with TEs from at least two different orders. *: p≤0.05. **B.** Biological functions of genes regulated by husk-specific enhancers overlapping MITEs and carrying potential TFBSs in enhancers. The bubbles represent the odds ratio measuring the enrichment in some Gene Ontology categories among genes potentially targeted by TFs binding the AAGGGATTTYTATTT 15-mer. **C.** Enrichment in TIR and LTR superfamilies among TFBSs located in V2-IST-specific enhancers. DTA: TIR *hAT*, DTC: TIR CACTA, DTM: TIR *Mutator*, RLC: LTR *Copia*, RLG: LTR *Gypsy*, RLX: LTR Unknown.

Most often, however, these MITEs did not overlap with any known TFBSs. Because the JASPAR database mostly contains TFBSs from *A. thaliana*, we hypothesized that this could be due to lack of a relevant maize TFBS motif in the database. Indeed, despite the identification of hundreds of maize TFs [36], JASPAR 2020 only contains 22 maize-specific binding motifs [37]. To investigate whether the detected MITEs contain a putative TBFS motif, we searched their sequences for conserved motifs by performing an enrichment analysis for 9- to 15-mers (see Materials and Methods). We found that MITEs included in husk-specific enhancers are enriched in 3 potential 15 pb TFBS motifs (Supplementary Table S8): AAATTAGTTYATTTT, AAGGGATTTYTATTT, and GTTCYCAAACTAGCC. By comparing these sequences to known plant TFBS motifs (JASPAR database), we found that they are most closely related to *Arabidopsis* HB-like, MYB-like, and MADS TFBS motifs, respectively. These motifs were significantly more present in MITEs of the *Pif/Harbinger* superfamily than in other MITEs superfamilies (odds ratio of 5.4, 9.5, and 9.0 respectively, see Fisher’s exact test results in Supplementary Table S9).

To get insights into the potential regulatory role of these enhancers, we investigated the biological functions of the genes targeted by enhancers carrying these motifs (Supplementary Table S10). Genes targeted by MITE-driven enhancers harboring the AAGGGATTTYTATTT motif are enriched for the biological process “lipid modification” and “leaf development” (see Figure 4B). Genes targeted by MITE-driven enhancers with the AAATTAGTTYATTTT and GTTCYCAAACTAGCC were enriched for “microtubule binding” and “isomerase activity” molecular functions, respectively.

V2-IST-specific enhancers were not enriched in particular TE superfamilies. However, when restricting the analysis to the TFBS parts of enhancers, we found that TFBSs from V2-IST specific enhancers were enriched for TIR transposon *Mutator* (TIR DTM—odds ratio of 12.9, resampled *χ*^2^ *p* = 0.05, Figure 4C). No other TIR and LTR superfamilies were significantly enriched in husk enhancer TFBS after *p*-value correction. The vast majority (70%) of TFBSs overlapping TIR transposon *Mutator* were from the AP2/ERF family. A gene ontology analysis revealed that candidate targets of enhancers carrying these TFBSs were enriched for biological processes related to nitrogen storage (GO:1901566, GO:0009073 and GO:0044283, see Supplementary Table S11).

## Discussion

In this study, we investigated the regulatory relationship between TFs and target genes in two leaf tissues at different developmental stages: V2-IST at seedling stage, and husk (bracts) at flowering. To do so, we used a bipartite network-based approach integrating several layers of information from heterogeneous data: epigenetic information (allowing to characterize an enhancer as active), genomic information (providing gene-candidate enhancer distance and annotation of TFBSs within enhancers) and transcriptomic information from 13 different tissues sampled at different developmental stages. This allowed us to provide functional insights into the regulatory role of enhancers that are activated in a tissue-specific manner.

An approach often used to analyze TFs-genes interactions is based on the inference of co-expression-based gene regulatory networks using machine learning. They have proven powerful to study maize regulatory networks of different tissues and to identify key regulatory TFs across development [38, 39, 40] or in response to the environment [41, 36]. However, they do not include information about enhancer sequences, and therefore do not allow to analyze the molecular origin of gene regulatory network modifications. In addition to identifying groups of co-regulated genes and corresponding key regulatory TFs involved in the regulation of leaf-specific functions at two different developmental stages, our methodology also allowed to identify the putative enhancers that regulate the target genes. This has two major advantages: (i) by characterizing the enhancers involved, it allows to investigate the molecular origin of tissue-specific regulation, and (ii) by identifying candidate enhancer-target genes pairs, it reduces the number of candidate target genes for each enhancer, thus limiting the number of candidates to be tested using molecular biology approaches.

Our study allowed us to identify regulatory modules corresponding to functions relevant to the tissue analyzed. For instance, in V2-IST, we find molecular functions expected to be found in an immature and growing tissue such as “cellular macromolecule biosynthetic processes”, “protein complex assembly” and “hormone response”. While we consolidate existing results, we also provide the key players (*i.e*., groups of co-regulated genes and their key regulatory TFs) involved in these functions. In addition to the to-be-expected modules, our approach also allowed us to discover regulatory modules of more unclear function, notably in husk. Husks are the bracts of the maize female inflorescence that provide a mechanical protection of the ear and growing silks (the styles of maize florets). In particular, husks ensure silks growth by protecting them from air evaporative demand and preserving their water status [42]. Its other biological functions, if any, remain poorly characterized. While the largest module, containing genes involved in the biosynthesis of cell walls, has been previously described in leaves in a late developmental stage [43], we identified a yet to be characterized husk-specific module. This module is regulated by two TFs, ERF039 and EREB10, whose functions are unknown in maize. Nevertheless, based on studies in other species, they are likely to be involved in response to biotic and abiotic stresses [44, 41]. For instance, an ortholog of ERF039 (also ortholog to *A. thaliana* ERF38) is involved in salt and osmotic tolerance in poplar [45]. Moreover, the AP2/EREB family, to which EREB10 belongs, participates in response to abiotic stress in soybean and maize [46, 47]. Their candidate targets include genes coding for TFs involved in the modification of photosynthesis and photomorphogenesis in response to abiotic stresses in many plant species [48, 49, 50, 51]. Husk growth is a component of the anthesis to silking interval, which is a good predictor of grain yield under stress [52]. Silk emergence out of the husk to be pollinated is indeed known to result from the balance between silk and husk growth rates, which are both responsive to abiotic constraints. Drought tolerance in maize thus partly relies on the coupling of tissue expansion in both vegetative and reproductive organs [53]. Our findings further support the importance for husk growth to be regulated in response to abiotic constraints, and pinpoint the genes and TFs involved in this process, thus allowing for their further molecular characterization. This will be particularly useful to understand how to improve maize response to drought.

In our attempt to characterize biological functions expressed in a tissue with limited biological characterization, we nevertheless encountered two limitations. First, the lack of functional annotation of maize genes in public databases prevented us to precisely annotate some of the husk-specific regulatory modules. Second, our TFBSs prediction in maize enhancers is based on motifs included in the JASPAR Plantae motifs database, which are mainly derived from ChIP-seq experiments performed in *Arabidopsis thaliana*. We are thus likely to be missing a number of tissue-specific functions regulated by maize-specific TFs. Despite these two caveats, we were able to provide candidate TFs and genes playing a key role in the expression of husk-specific biological functions. Our approach, coupled with the rapidly increasing data available on maize-specific TFBS motifs [54], will thus improve our capacity to concomitantly identify enhancers, TFs and genes involved in the regulation of the expression of biological functions in poorly characterized tissues.

Our results support the role of TEs as functional actors in the tissue-specific regulation of biological functions involved in leaf differentiation. We find that a substantial amount (~ 60%) of the enhancers analyzed include TE sequences. This is higher than estimated in previous studies [8, 10] and likely arises from the fact that we used an upgraded maize TE database [35], which allowed a more in depth characterization of TEs within enhancers. Several studies have shown that TEs can modify gene regulation under stress conditions, for instance in rice [55] and in maize [21]. But the underlying mechanisms are still unclear. Several cases of TE-driven enhancers involved in the regulation of specific genes have been described in plants. For instance, a *Hopscotch* LTR retrotransposon regulates the domestication gene *tb1* in maize through a long-range interaction [56, 4]. In *A. thaliana*, LINE *EPCOT3* is involved in the neo-functionalization of the *Cyp82c2* gene, thus contributing to chemical diversity and pathogen defense [17]. *In Brassica napus*, a CACTA transposon acts as an enhancer to stimulate expression of the *BnaA9.CYP78A9* gene and silique elongation [19]. In maize, a recent analysis of chromatin accessibility at the genome-wide level revealed that TEs, in particular LTR retrotransposons, contribute to gene regulation as *cis*-regulatory elements [10]. But this study did not connect TE-driven enhancers to their target genes.

Here, by taking advantage of the concomitant characterization of enhancers and their target genes, we discovered the TE sequences associated with the enhancers, but also the functions that they regulate. This allowed us to show that TIR transposons and MITEs can directly regulate gene expression through their domestication as enhancers in maize, and are involved in the regulation of tissue-specificity. Interestingly, while Zhao *et al*. (2018) pointed mainly to the role of LTR retrotransposons [10], we point here to the role of TIR transposons and MITEs in tissue-specific regulation, suggesting that these elements may be involved in tissue-specificity. Moreover, we show that two distinct TE families, TIR transposon *Mutator* and MITE *Pif/Harbinger*, have provided TFBSs to enhancers regulating the expression of genes from two distinct pathways: nitrogen storage in V2-IST, and late-stage leaf development in husk, respectively. This highlights potential selection of different families to rewire regulatory networks across development.

Finally, through analysis of MITE *Pif/Harbinger*-driven enhancers, we discovered a new potential TFBS motif involved in the regulation of husk development, which is likely recognized by a MYB-like TF. MYB-like TFs have been shown to be highly expressed in the late stage of maize leaf development, and to play a role in the regulation of circadian rhythm and photosynthesis regulation [57]. Here, we propose that part of the gene regulatory network underlying late-stage husk development has been shaped by the domestication of MITE elements carrying MYB-like TFBSs. Our results complete previous ones showing that the transposition of MITEs have helped amplify specific TFBS and rewire the gene regulatory networks controlling key biological processes in several species, including the response to stress and flowering time in peach and other *Prunus*, and fruit rippening in tomato [58]. This highlights the power of our methodology to identify potential new TFBSs.

To conclude, our combined analysis of maize genomic, epigenomic and transcriptomic data using bipartite networks allowed us to analyze the role of enhancers in the development of leaf tissues. We were able to identify key actors involved in leaf development at different molecular levels, from the biological functions involved, to the underlying enhancer-target gene pairs and key transcription factors. We highlighted the role of TIR transposable elements as important actors of tissue-specific gene regulatory expression wiring, through their domestication as distal *cis*-regulatory sequences. We also discovered new potential TE-based TFBSs. By connecting enhancers to their target genes, and identifying the biological functions they potentially regulate, our work opens new avenues to study the impact of enhancer structural variation on the wiring of gene regulatory networks and, ultimately, to the underlying phenotype.

## Materials and Methods

### Previously generated data

We use the coordinates of active enhancers and corresponding mRNA-seq data from two different tissues, husk (the soft inner leaves surrounding the ear) and V2-IST (the inner stem of stage V2 seedlings) from the B73 maize line. These data were generated in a former work [8]. Briefly, active enhancer coordinates were obtained by intersecting DNAse I hypersensitivity DNAse-seq, histone mark H3K9ac ChIP-seq and DNA methylation bisulfite-sequencing profiles (see [8]). RNA-seq data for six replicates for both husk and V2-IST were also provided by Oka and colleagues (raw fastq files with 100 bp single-end reads). More information about plant growth, RNA extraction and library preparation can be found in [8].

### Generation of mRNA-seq data: mRNA extraction, library preparation and sequencing

We generated 150 bases paired-end mRNA-seq data from 11 tissues. Tissue types, growing conditions and sampling are summarized in Supplementary Table S2. For all tissues, mRNAs were extracted from three independent plants (biological triplicate), except for hypocotyl and roots, where a replicate is a pool of three different plants and for 17DAP and 35DAP seeds, where a replicate is a pool of seeds from a single ear. For leaf, internode, silk, tassel, immature ear and 17DAP seed, RNAs were isolated with with Trizol (Invitrogen ref.15596018) and β-mercaptoethanol (SIGMA ref. M3148-25ML) reagents. Supernatant was recovered and RNA purified using Qiagen RNeasy Plant Mini kit (ref. 74904) following manufacturer’s instructions. Then, a Qiagen RNAse-free DNAse set (ref. 79254) was applied to remove the residual DNA. A different protocol was used for 35DAP seed mRNA extraction: RNAs were extracted with 4.5 ml of buffer (10 mM Tris-HCl, pH7.4, 1 mM EDTA, 0.1 M NaCI, 1% Sodium Dodecyl Sulfate) and 3 ml of phenol — chloroform — isoamyl alcohol mixture 25:24:1. The supernatant was extracted one more time with the same phenol solution in order to eliminate proteins and starch. The nucleic acids were precipitated by addition of 0.1 vol of 3M sodium acetate pH5.2 and 2 vol of 100% ethanol. After precipitation RNA were rinsed one time with 70% ethanol and the pellets dissolved in RNase-free water. Purification was done with a DNAse treatment RNase-Free DNase Set (Qiagen, Hilden, Allemagne) and then RNeasy MinElute Cleanup Kit (Qiagen, Hilden, Allemagne). Quality of total RNA samples was assessed using the Agilent 2100 bioanalyzer (California, USA) according to manufacturer’s recommendations. Library construction was generated by the IPS2-POPS platform. Briefly, mRNAs were polyA selected, fragmented to 260 bases and libraries were built using the TruSeq stranded mRNA kit (Illumina^®^, California, U.S.A.) with an Applied BioSystem 2720 Thermal Cycler and barcoded adaptors. Barcoded libraries were sequenced on Illumina HiSeq4000 at Genoscope, in paired-end (PE) with 150 bases read length. Approximately 20 millions of paired-end reads were produced for each sample.

### RNA-seq data pre-processing

Raw illumina reads were aligned to the AGPv4 version of the B73 maize genome using STAR [59] and the following options: —runMode alignReads —alignIntronMin 5 —alignIntronMax 60000 —runThreadN 32 —readFilesCommand gunzip -c —quantMode GeneCounts SortedByCoordinate —outSAMprimaryFlag AllBestScore —outFilterMultimapScoreRange 0 —outFilterMultimapNmax 20 —alignEndsType Local —sjdbGTFtagExonParentGene gene_id. Read counts per gene were calculated with STAR from reads unambiguously mapped on genes. With these settings, over all tissues, 91.1% of reads (median) were mapped unambiguously to a gene for the Oka *et al*. dataset (ranging from 87.9% to 92.1%), and 95.8% for the AMAIZING Gene Atlas dataset (ranging from 92.8% to 97.7%).

Counts per gene from both datasets were then pooled and normalized together using the tissue-aware smooth quantile normalization from the R bioconductor *YARN* package version 1.1.1 [60], using the normalizeTissueAware function with the *method* = *”qsmooth”* option. Data were then corrected for single-end/paired-end batch effect using the removeBatchEffect function from the R bioconductor *limma* package version 3.28.2.

### Enhancer definition and TFBS identification

Candidate enhancer sequences were extracted from the bed files containing coordinates of enhancers from [8] and the AGPv4 maize genome sequence using bedtools getfasta. They were scanned for Transcription Factor Binding Sites (TFBS) using the *FIMO* software from the *MEME* v. 5.0.5 suite [61] using default parameters. To this end, we retrieved the MEME core plants position frequency matrix files corresponding to the binding sites of 489 transcription factors available in JASPAR database (accession october 31 2019) [37]. Matches with a q-value (Benjamini-Hochberg corrected *p*-value) lower than 0.01 were retained.

Significance of the TFBS results was tested by comparing the number of bases covered by TFBSs in original candidate enhancer sequences to this of random sequences obtained by shuffling enhancers dinucleotides. To this end, for each enhancer, we generated 1000 random sequences using the BiasAway software [62] and computed resampling *p*-values by counting the number of random sequences for which TFBS coverage exceeded the one of the original enhancer.

We compared the TFBS motif content of husk and V2-IST enhancers by using the AME software from the *MEME* suite version 5.0.5 [63]. Enrichment in particular TFBSs among husk enhancers was estimated by setting husk as primary and V2-IST as background sequences. This procedure was swapped to obtain V2-IST enhancer TFBS enrichment. We tested enrichment for motifs using the Fisher exact test, and *p*-values were corrected for multiple testing using the Bonferroni method. *E*-value threshold was set to default (*E* ≤ 10).

### TF-gene network building

We built tissue-specific regulatory networks using the PANDA and LIONESS softwares [22, 25]. PANDA represents regulatory relationships between transcription factors and genes as a bipartite network, nodes being either transcription factors or genes, and edge weight being proportional to the strength of the TF-gene regulatory relationship. The method requires as input a prior representing potential regulatory relationships between transcription factors and genes. The prior is a gene × transcription factors matrix of zeros and ones, were ones indicate the presence of a putative transcription factor binding site in a *cis* -regulatory region of the gene, and zero its absence. This prior edge matrix is then updated using a protein-protein interaction (PPI) matrix that represents interactions between transcription factors and a gene co-expression matrix. This relies on a message-passing algorithm that verifies both the “responsibility” and the “availability” of each edge [22]. The final PANDA output is an “aggregate” network model representing gene regulation in a specific dataset. LIONESS is a mathematical framework that extracts networks for individual samples from such an aggregate regulatory network [25].

In this study, the prior TF-gene interaction matrix was obtained by crossing enhancer coordinates with gene coordinates. All enhancers within 250 kb upstream or downstream of a gene transcription start site were annotated as a potential regulator, and prior edges between this gene and each transcription factor mapping to those enhancers were set to 1. All other prior edges were set to 0. The co-expression matrix was obtained from the mRNA-seq normalized data from the 45 samples (13 tissues). In absence of a detailed protein-protein interaction matrix for plants, we used an identity matrix. Using PANDA and LIONESS, we generated 46 networks: a global network (PANDA), and 45 sample-specific ones (LIONESS). Raw sample-specific edges weights (*EW*) were log-transformed (*logEW*) using the following formula: for each sample *i* and edge *e*:

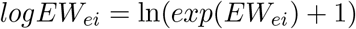

Edges that were set to 0 in the prior were set to 0 in the all the following analyses.

Enhancer-gene TSS distances thus obtained were also used to compute the distributions of distances between enhancers and nearest gene TSS. Briefly, the closest gene was the one whose TSS was the closest from the gene in absolute value.

### Identification of most likely target gene

In order to identify the most likely target gene of each enhancer, we computed the average edge weight for each enhancer-potential target gene pair. For each pair of enhancer *e* and potential target gene *g*, the average edge weight (*avgEW_g_*) was computed as the average of the edges (logEW) from each TF *t* to the gene. Only TFs with a potential factor binding site in the enhancer were included. With *T_e_* the number of TFBSs in the enhancer:

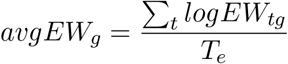

### Tissue-specific gene targeting

To identify genes that were differentially targeted between husk and V2-IST tissues, edges weights were compared between tissues-conditions using a linear regression performed with the R bioconductor *limma* R package version 3.28.2.

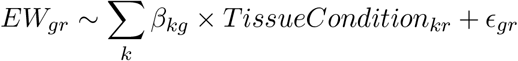

with *k* being the tissue-condition and *r* the replicate.

Genes that were targeted by edges with Benjamini-Hochberg corrected *p*-values under 0.01 were considered as differentially targeted.

### Identification of tissue-specific TF-genes regulatory modules

To identify tissue-specific regulatory modules, we first build one husk-specific and one V2-IST-specific network by averaging the LIONESS sample-specific networks across the replicates for each tissue. We then identified the TF and genes that changed the modularity of the networks between V2-IST and husk by running ALPACA [26], setting in turn the V2-IST and husk networks as background. This function outputs one list of regulatory modules—*i.e*. groups of TFs that regulate groups of genes based on the edge weights—for each tissue-specific network. We then compared the gene content of the regulatory modules between husk and V2-IST by computing pairwise jaccard indexes. The maximum jaccard index was conserved for each module in each tissue. Modules with maximum jaccard index over 0.5 were annotated as shared between tissues, and modules with maximum jaccard index under 0.5 were annotated as tissue-specific.

### Gene Ontology enrichment analyses

We performed Gene Ontology enrichment analyses using the R bioconductor topGO package [64], using the Fisher test and the elim method. As the tests are not independent, no multiple testing correction can be applied. Instead, following the guidelines from the users’ manual, we filtered uncorrected *p*-values with a stringent threshold of 0.01. For GO enrichment analysis of genes in tissue-specific modules, if the module contained more than 100 genes, we performed gene enrichment analyses on the 100 genes that were the most connected to TFs within the regulatory module (top differential modularity genes from the ALPACA results). The gene ontology database used in this analysis was generated by combining publicly available annotations [65] obtained from InterproScan5, Arabidopsis and uniprot from https://datacommons.cyverse.org/browse/iplant/home/shared/commons_repo/curated/Carolyn_Lawrence-Dill_maize-GAMER_maize.B73_RefGen_v4_Zm00001d.2_0ct_2017.r1/d.non_red_gaf (January 2020), and removing any redundancy.

### Annotation of transposable elements in enhancers and TFBS

We annotated the transposable elements of the B73 genome (AGPv4) using REPEATMASKER v4.0 (http://www.repeatmasker.org) with the updated MTEC database provided by Ou and colleagues [35, 66, 67]. We annotated enhancers and TFBSs using this database and the data.table R package [68].

We tested for enrichment in a particular transposable element superfamily among husk-specific or V2-IST-specific enhancers using a *χ*^2^-test. In order to take into account the genomic location of the enhancers in the TE enrichment analysis, we computed a null distribution of *χ*^2^ values using 1,000 resamplings of genomic sequences with same length, chromosome and distance to nearest gene TSS as the original list of enhancers used.

### Motif discovery in MITE sequences

We searched MITE sequences overlapping husk-specific enhancers for motifs using the MEME software [69]. Because TFBSs from plants are typically 11 nt long, we searched for motifs of length 9 to 15 nt. We filtered out motifs with an E-value over 10^−4^. We then used the online version of Tomtom [70] to compare these motifs with known TFBS motifs available in the JASPAR 2018 non-redundant core plants database. We filtered out all motifs with a *p*-value greater than 0.01.

## Acknowledgement

This work was supported by the Investment for the Future ANR-10-BTBR-01-01 Amaizing program, and we thank Alain Charcosset for its coordination. We thank Harry Belcram and Agnes Rousselet for help in tissue grinding and extractions, Peter Rogowsky for hypocotyl and root experiment coordination, Jerome Laplaige for root and hypotocyle sampling and extraction, Sylvie Coursol for internode experiment coordination, Sandrine Chaignon for internode sampling and extractions, and Carine Palaffre for grain experiment set up and sample collection. The GQE and IPS2 laboratories and the POPS platform benefit from the support of Saclay Plant Sciences-SPS (ANR-17-EUR-0007). The POPS platform benefits from the privileged access to the Genoscope sequencing facility. We thank the Genotoul bioinformatics platform Toulouse Midi-Pyrenees for providing computing and storage resources, Shujun Ou for assistance with the EDTA program that we used for TE annotation, Anthony Mathelier for help with TFBS identification, and Megha Padi for assistance with ALPACA.

## Authors Contribution

C.V., M.F. and M.S. conceived the experiments. C.V. and J.J. secured funding. C.V. and M.F. performed the experiments and analyzed data. O.T. coordinated greenhouse culture, sampling strategy and sample collection. A.V., C.V., J.J. and O.T collected samples. A.V. performed tissue grinding and mRNA extractions. S.P. generated mRNA-seq libraries. J.R. aligned mRNA-seq reads. M.F. and C.V. wrote the article. J.J., M.L.K., M.S. and O.T. provided expertise, feedback and corrected the manuscript.

## Data availability

All steps of the experiment, from growth conditions to bioinformatic analyses, were managed in CATdb database [71] (http://tools.ips2.u-psud.fr/CATdb/) with Project ID NGS2017_06_AMAIZING, experiment Amaizing_B73_WW. This project has been submitted from CATdb into the international repository GEO (Gene Expression Omnibus, Edgard R. et al. 2002, http://www.ncbi.nlm.nih.gov/geo) with ProjetID GSE151455. These data are available upon request to Clementine Vitte, and will be made publicly available upon manuscript acceptance.

---

**Figure S1:**
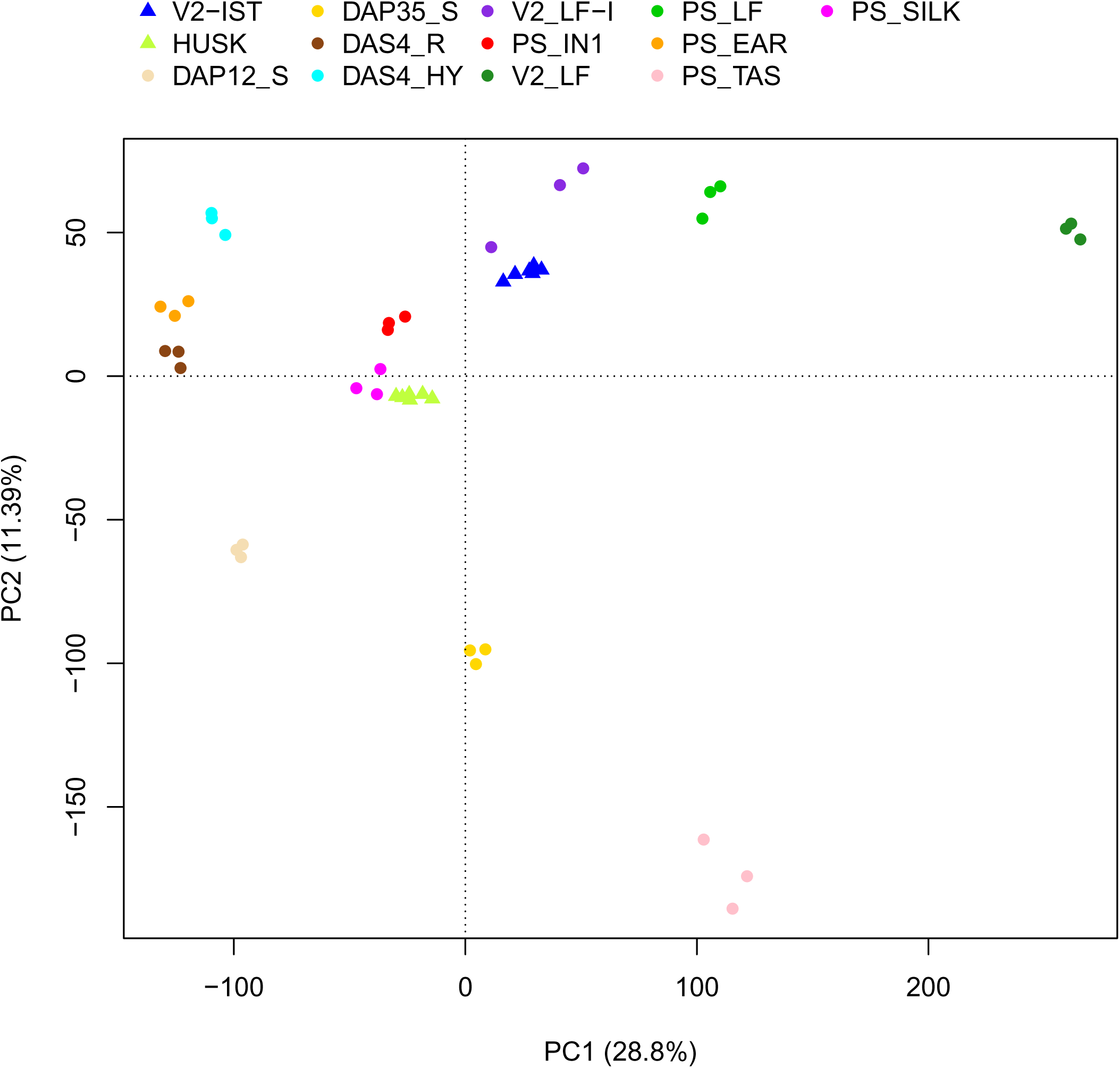
Principal component analysis of normalized and batch-corrected RNA-seq expression data from the GeneAtlas AMAIZING dataset (circles) and Oka *et al*. dataset (triangles).

**Figure S2:**
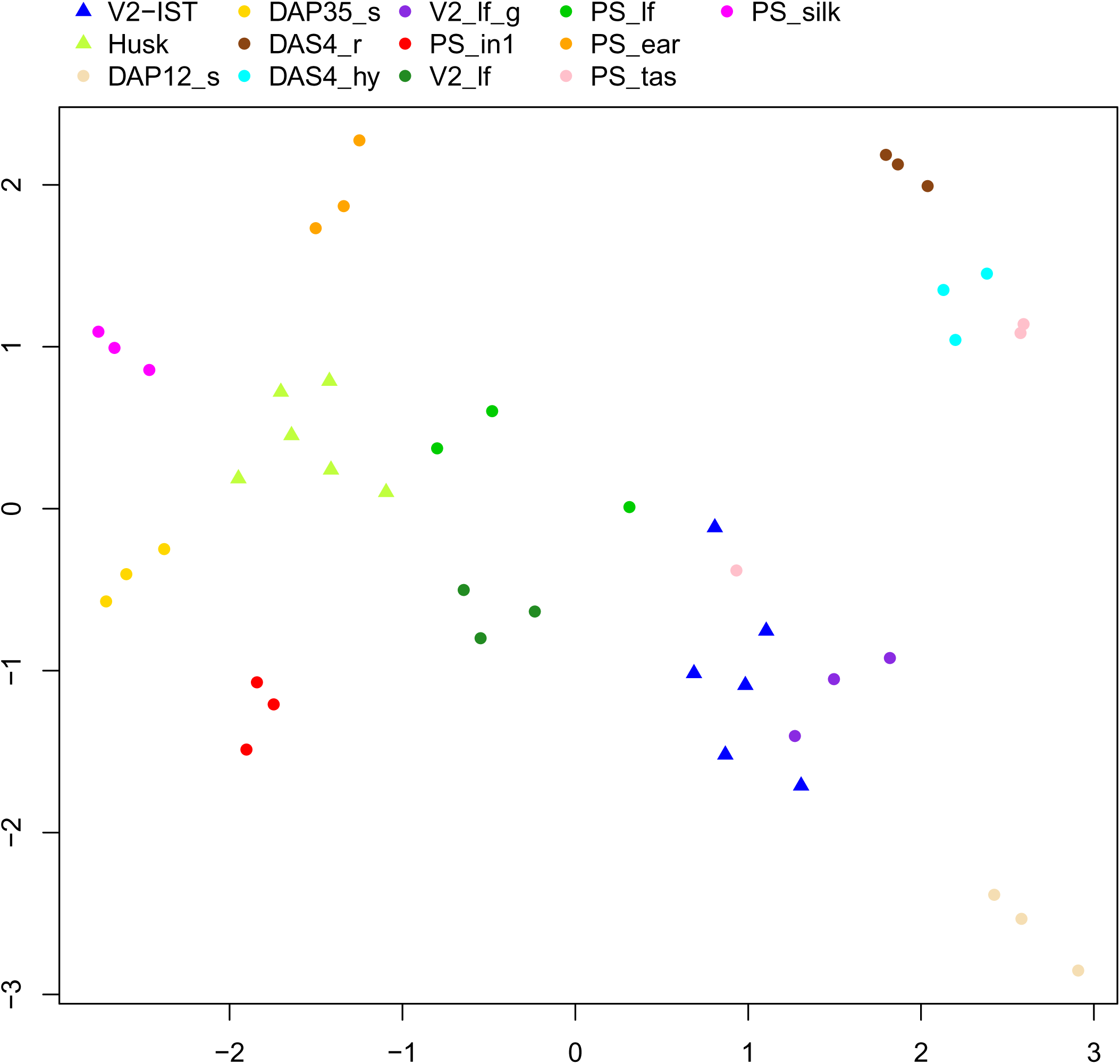
UMAP representation of edge weights from sample-specific networks. Circles correspond to samples from the GeneAtlas AMAIZING dataset and triangles to the Oka *et al*. dataset.

**Figure S3:**
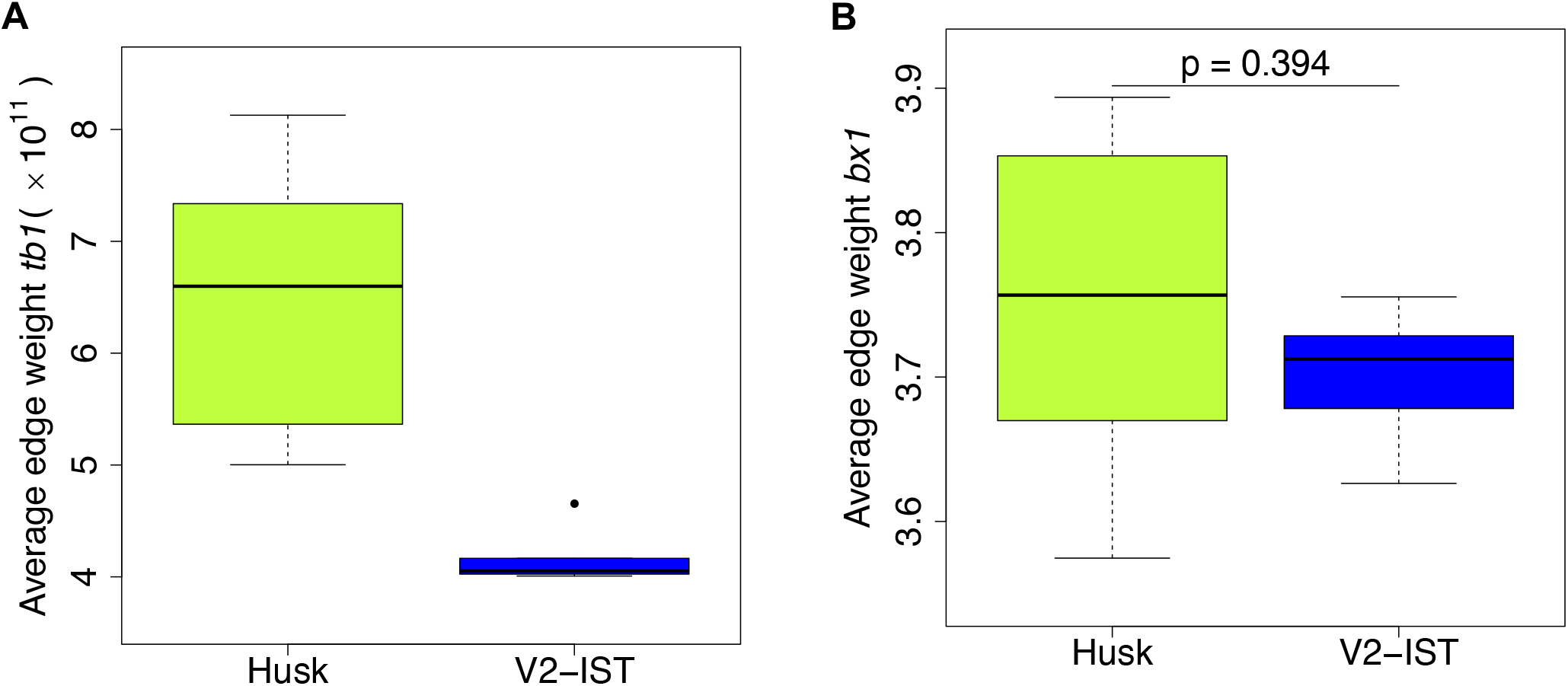
Average edge weights between known enhancers and their target genes. **A.** Enhancer of *tb1*, which is active in husk but not in V2-IST: distribution of average edge weight between *tb1* and the transcription factors that have binding sites in its enhancer for husk and V2-IST samples. **B.** Enhancer of *bx1* (DICE), which is found to be active in both husk and V2-IST: distribution of the average edge weight between *bx1* and the transcription factors that have binding sites in DICE for husk and V2-IST samples.

**Figure S4:**
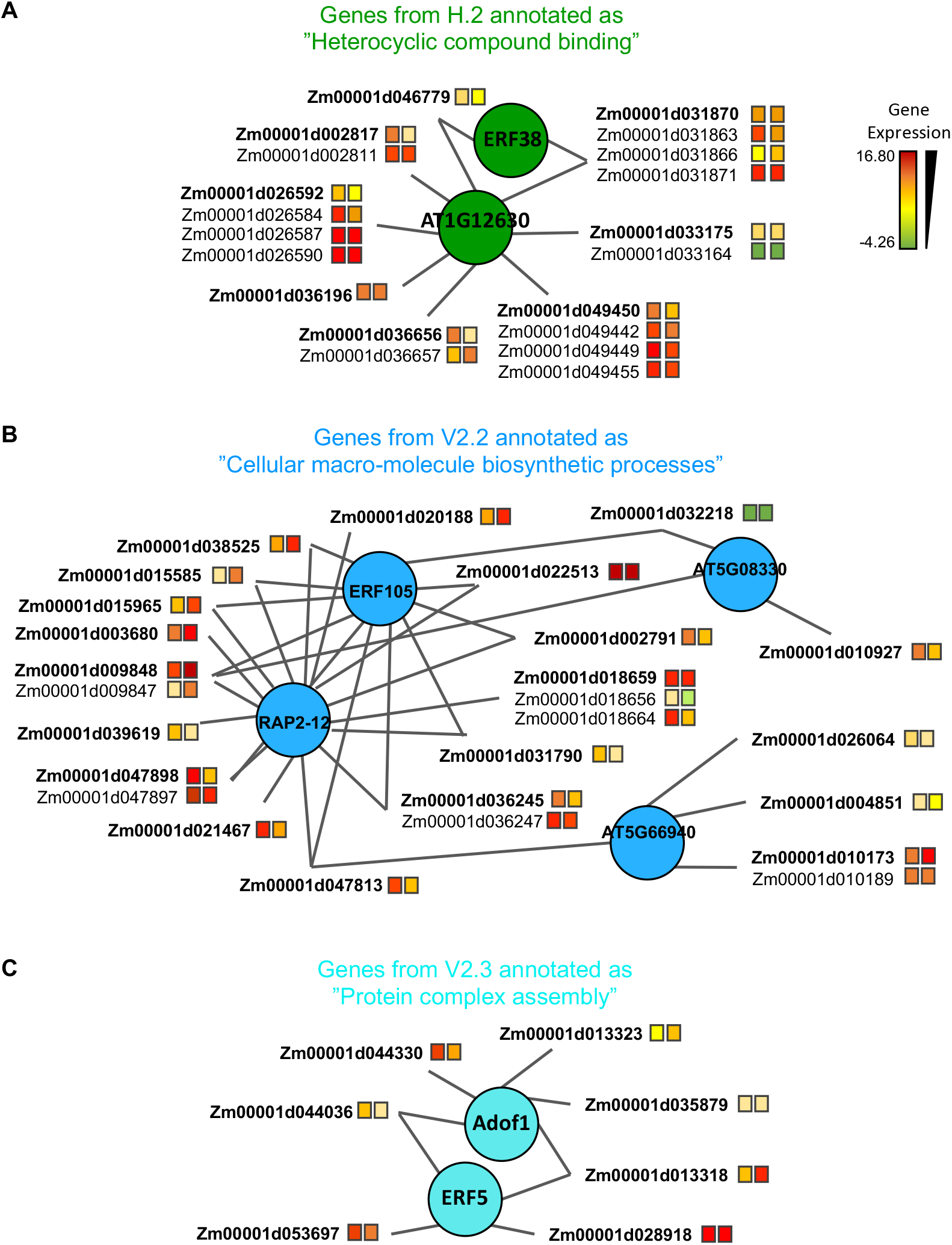
Detailed view of the husk and V2-IST-specific modules. textbfA. Detailed view of the part of the husk-specific H.2 module that shows the genes annotated as “heterocyclic compound binding”. **B.** Detailed view of the part of the V2-IST-specific V2.2 module that shows the genes annotated as “cellular macromolecule biosynthetic processes”, showing the connection of all the genes involved in this biological process. **C.** Detailed view of the part of the V2.3 module that shows the genes annotated as “protein complex assembly”, showing the connection of all the genes involved in this biological process. **A-C.** The color of the squares beside the names represent the genes average expression levels in husk (left square) and V2-IST (right square). When several genes are potentially targeted by the same enhancer, they are represented with a common edge, and the top target is ranked first and highlighted in bold. The TFs that regulate the genes are represented as circles. Because TFBS annotation arise from *Arabidopsis thaliana*, names of TFs are these of this species. Maize orthologs of *erf38, At1g12630, rap2-12 erf5* and *adof1* are *erf039, ereb10, bbr4, ereb210, ereb61*, and *dof7* respectively. Similar information can be retrieved for all genes of the module using the R application we developped https://maud-fagny.shinyapps.io/TF-gene_network_Maize/.

**Figure S5:**
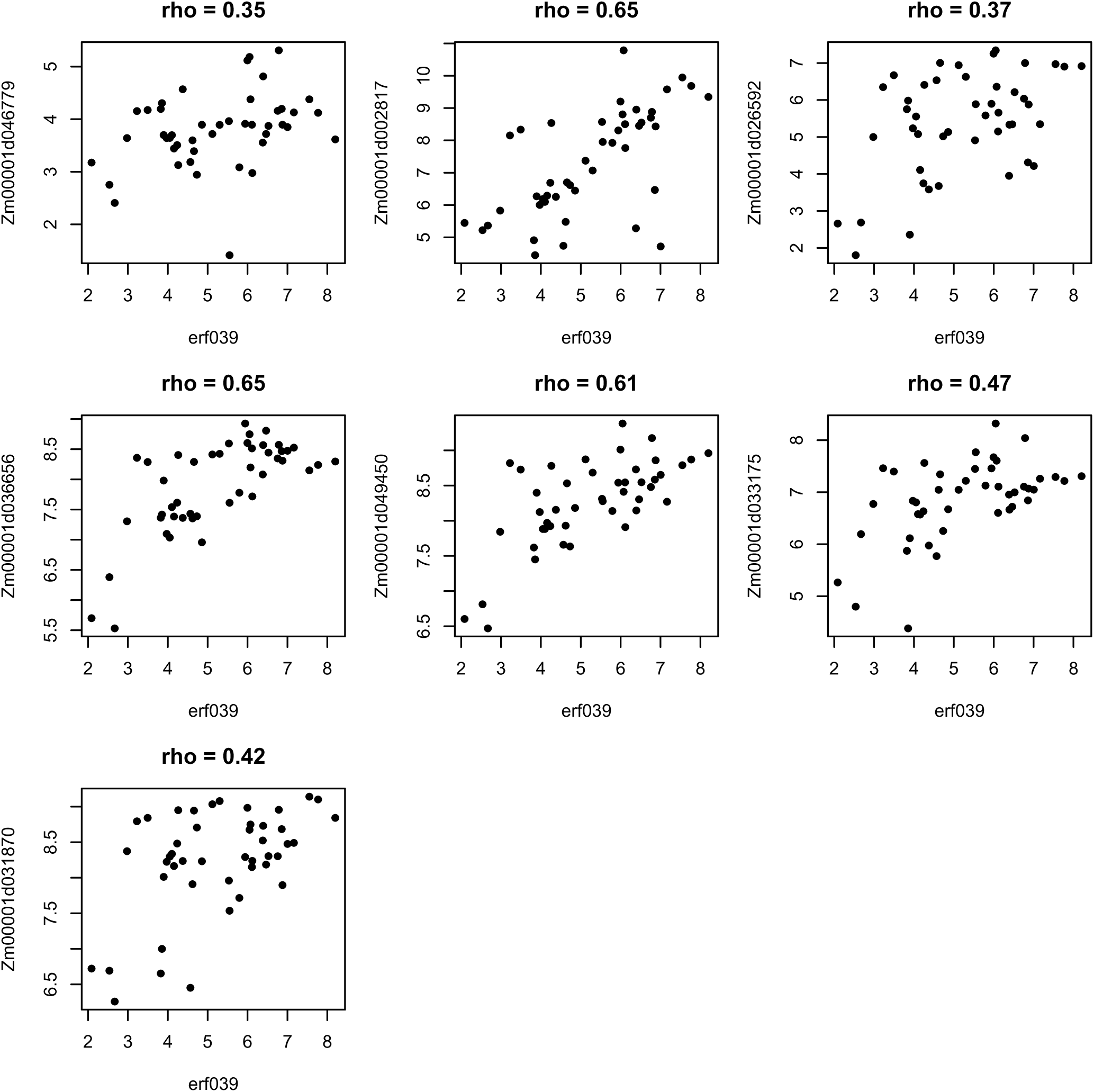
Correlations of expression levels between *erf039*, the maize ortholog of *erf38* and ERF38 target genes involved in heterocyclic compound binding across all 45 samples. Only the top target gene for each ERF38-binding enhancer is represented. Each panel represents the correlation between *erf039* (x-axis) and one of its target gene annotated as “heterocyclic compound binding” (y-axis). Spearman’s rho values are indicated on top of each graph.

**Figure S6:**
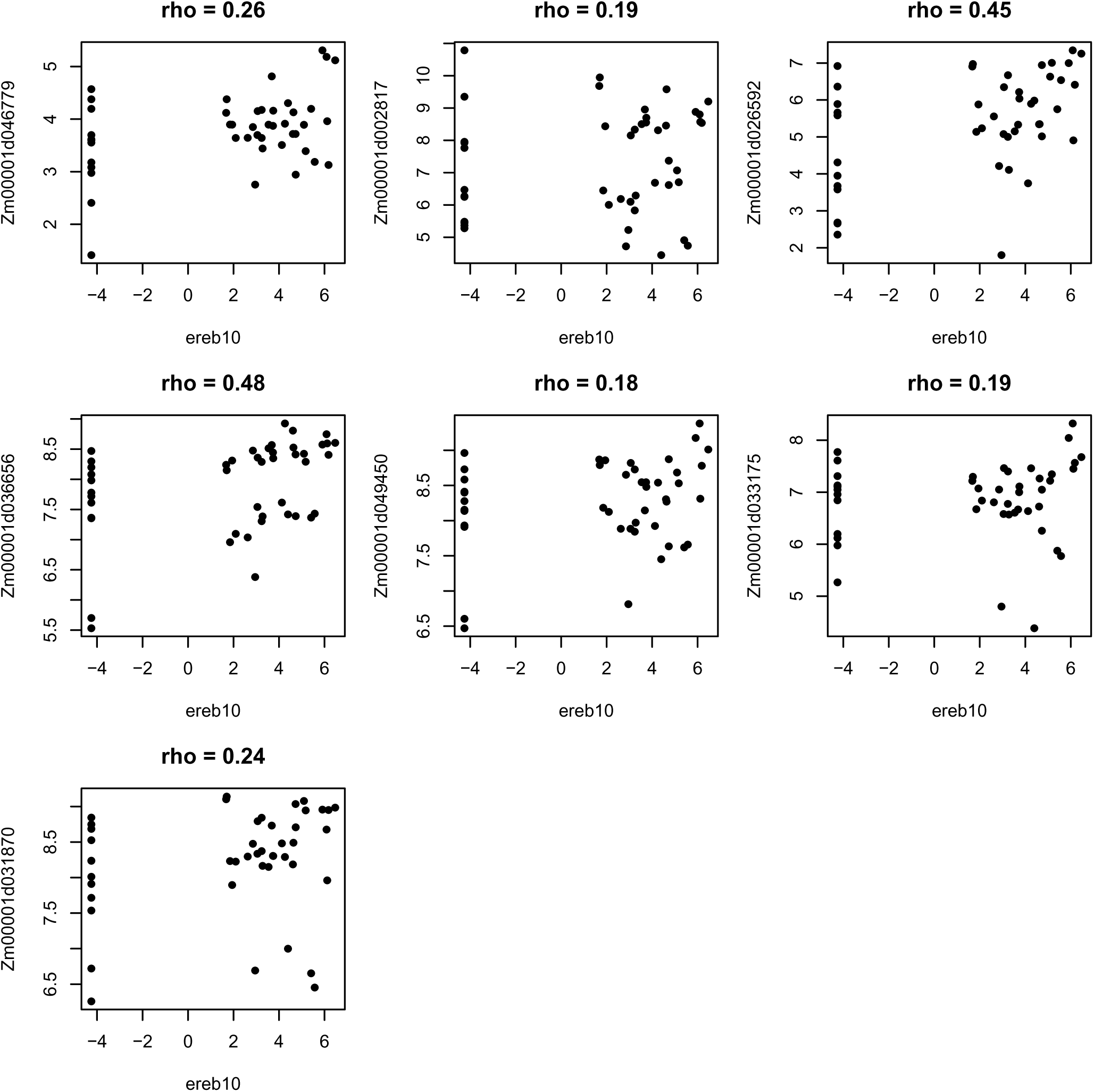
Correlations of expression levels between *ereb10*, the maize ortholog of *At1g12630* and AT1G12630 target genes involved in heterocyclic compound binding across all 45 samples. Only the top target gene for each AT1G12630-binding enhancer is represented. Each panel represents the correlation between *ereb10* (x-axis) and one of its target genes annotated as “heterocyclic compound binding” (y-axis). Spearman’s rho values are indicated on top of each graph.

**Figure S7:**
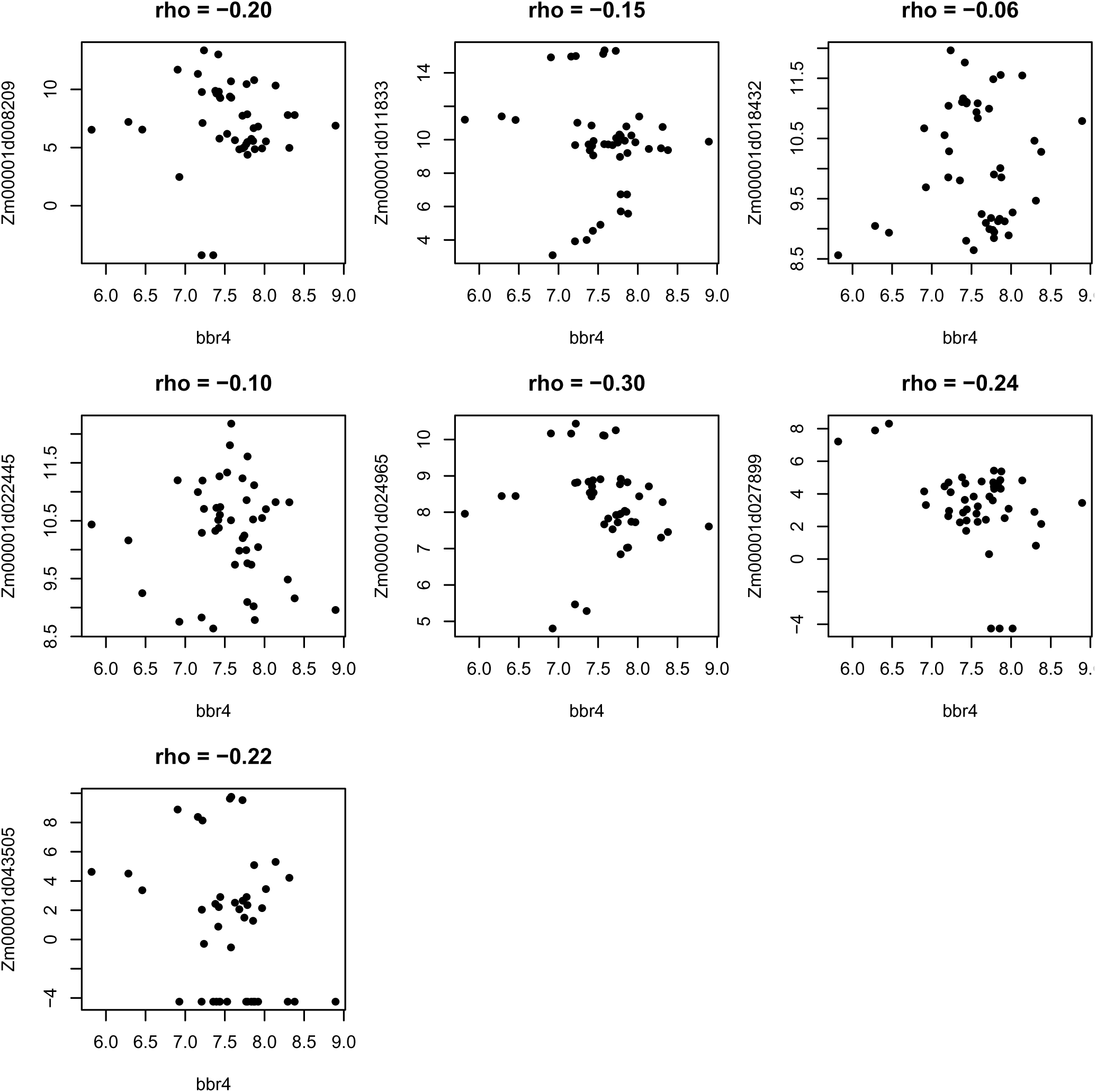
Correlations of expression levels between *bbr4*, the maize ortholog of *bpc5*, and BPC5 target genes involved in hormone response across all 45 samples. Only the top target gene for each BPC5-binding enhancer is represented. Each panel represents the correlation between *bbr4* (x-axis) and one of its target genes annotated as “hormone response” (y-axis). Spearman’s rho values are indicated on top of each graph.

## Supplementary Tables

**Table S1:** Supplementary_Dataset_S1_TFBS_enrichment_enhancers.xlsx. Binding sites mapping preferentially in husk or V2-IST enhancers.

**Table S2:** Supplementary_Dataset_S2_Nomenclature_samples.xlsx. Samples characteristics.

**Table S3:** Supplementary_Dataset_S3_Gene_ontology_enrichment_Husk.xlsx. Gene Ontology enrichment analysis of genes preferentially targeted in husk.

**Table S4:** Supplementary_Dataset_S4_Gene_ontology_enrichment_V2-IST.xlsx. Gene Ontology enrichment analysis of genes preferentially targeted in V2-IST.

**Table S5:** Supplementary_Dataset_S5_jaccard_mdex_communities.xlsx. Comparison of the gene content of husk and V2-IST regulatory modules using jaccard index.

**Table S6:** Supplementary_Dataset_S6_Shared_modules_GO.xlsx. Gene Ontology enrichment analysis of shared modules between husk and V2-IST tissues.

**Table S7:** Supplementary_Dataset_S7_Tissue_Specific_modules_GO.xlsx. Gene Ontology enrichment analysis of tissue-specific modules for husk and V2-IST networks.

**Table S8:** Supplementary_Dataset_S8_Husk_MITE_TFBS_similarity.xlsx. Motif enrichment analysis in MITEs overlapping husk-specific enhancers.

**Table S9:** Supplementary_Dataset_S9_MITEs_motifs.xlsx. Enrichment analysis of MITE putative TFBS motifs among *Pif/Harbinger* elements.

**Table S10:** Supplementary_Dataset_S10_Husk_MITE_target_GO.xlsx. Gene Ontology enrichment analysis of genes targeted by MITE-containing husk-specific enhancers carrying each of the 3 motifs.

**Table S11:** Supplementary_Dataset_S11_V2-IST_TIR_target_GO.xlsx. Gene Ontology enrichment analysis of enhancers carrying AP2/ERF transcription factors overlapping with TIR *Mutator* TE.

